# Invisible noise obscures visible signal in insect motion detection

**DOI:** 10.1101/098459

**Authors:** Ghaith Tarawneh, Vivek Nityananda, Ronny Rosner, Steven Errington, William Herbert, Bruce G. Cumming, Jenny C. A. Read, Ignacio Serrano-Pedraza

**Author notes:** **Contributions:** - Designed research: GT, VN, RR, JR, ISP - Performed research: GT, SE, WH, JR, ISP - Contributed unpublished reagents/analytic tools: GT, ISP - Analyzed data: GT, JR, ISP - Wrote the paper: GT, JR, ISP, BGC.

## Abstract

The motion energy model is the standard account of motion detection in animals from beetles to humans. Despite this common basis, we show here that a difference in the early stages of visual processing between mammals and insects leads this model to make radically different behavioural predictions. In insects, early filtering is spatially lowpass, which makes the surprising prediction that motion detection can be impaired by “invisible” noise, i.e. noise at a spatial frequency that elicits no response when presented on its own as a signal. We confirm this prediction using the optomotor response of praying mantis *Sphodromantis lineola*. This does not occur in mammals, where spatially bandpass early filtering means that linear systems techniques, such as deriving channel sensitivity from masking functions, remain approximately valid. Counter-intuitive effects such as masking by invisible noise may occur in neural circuits wherever a nonlinearity is followed by a difference operation.

## 2 Introduction

Linear system analysis, first introduced in visual neuroscience decades ago (Campbell and Robson, 1968; Carandini, 2006), has been highly influential and continues to be successfully applied in several domains including contrast, disparity and motion perception (Anderson and Burr, 1985; Batista et al., 2013; Burge and Geisler, 2014; Carandini et al., 2005). This is despite the fact that neurons have many well-known nonlinearities. For example, nonlinearity is fundamental in accounting for our ability to perceive the direction of moving patterns (Adelson and Bergen, 1985; Clifford and Ibbotson, 2002; Emerson et al., 1992). Increasingly, contemporary models in neuroscience consist of a cascade of linear-nonlinear interactions (Chichilnisky, 2001; Hunter and Korenberg, 1986; Meister and Berry, 1999). At each stage of these models, inputs are pooled linearly and then processed with a nonlinear operator such as divisive normalization. It is therefore somewhat surprising that linear systems analyses work as well as they do.

In a linear system, noise injected at a frequency to which a sensory system does not respond has no effect on the system’s ability to detect a signal. This property is often taken for granted in the study of perception. For example, since humans cannot hear ultrasound, our ability to discriminate speech is not affected by the presence of ultrasound noise. In fact, our auditory system consists of frequency-selective and independently-operating linear channels so that even noise at frequencies to which we are sensitive may not affect our hearing performance, if the noise is not detected by the same channel as the signal (Patterson and Nimmo-Smith, 1980). Similar frequency-selective channels mediate the detection of contrast and motion in the visual system (Anderson and Burr, 1985; Blakemore and Campbell, 1969; Campbell and Robson, 1968; Graham and Nachmias, 1971; Sachs et al., 1971). Whether a system consists of multiple channels or not, it may seem obvious that the performance of the system cannot be affected by noise at frequencies to which the system does not respond. However, this property does not necessarily hold in nonlinear systems. It is possible to build a nonlinear system that is unresponsive to signals at a particular frequency but whose performance is significantly affected if noise of that frequency is added to a signal. Although there are many well-understood nonlinearities in vision (Badcock, 1984; Burr, 1980; Burton, 1973; Chen et al., 1993; Lawton, 1984; Marr and Hildreth, 1980; Morrone and Burr, 1988; Pollen et al., 1988; Zhou and Baker, 1993), interactions of this kind are often ignored (Harvey and Gervais, 1978; Legge, 1976; Maffei and Fiorentini, 1973; Stromeyer and Julesz, 1972). The prominent models in the domain of motion perception, for example, include well-known nonlinearities but are still assumed to not respond to noise outside their frequency sensitivity band (Anderson and Burr, 1989).

Here, we show that this assumption is not generally true for the standard models of motion perception. The nonlinearity of these models means that a moving signal at a highly visible frequency can be ”‘masked”’ (made less detectable) by noise at frequencies outside the detector’s sensitivity band (i.e. *invisible noise*). So far, this effect has been neglected because it does not occur when the filtering prior to motion detection is spatially bandpass, as it is in mammals. Masking techniques in humans and other mammals could therefore be applied successfully while ignoring this effect. However, in insects, early filtering is lowpass, and so we predict that invisible noise will be able to obscure a moving signal.

To test this prediction, we used the optomotor response of the praying mantis *Sphodromantis lineola*, in response to drifting gratings with and without noise. We have previously shown that the optomotor response is most sensitive to gratings at around 0.03 cycles per degree (cpd) and is largely insensitive to signals below 10^−2^ cpd (Nityananda et al., 2015). However, we show here that noise as low as 10^−3^ cpd – an order of magnitude lower – has the same effect as noise of the same amplitude presented at the optimal spatial frequency. This is quite different from published results in humans (Anderson and Burr, 1985), where noise has most effect when presented at spatial frequencies close to the optimal frequency of the relevant channel, and has no effect when presented at frequencies to which the organism is not sensitive. However, we show that the same model structure correctly predicts the qualitatively different behaviour in the two species, reflecting a difference in early filtering. Thus a profound difference between the behaviour of insects and of humans actually helps to confirm that both species use a similar mechanism to compute visual motion.

## 3 Results

### 3.1 Modelling biological motion detection

The standard models of early biological motion detection are the Hassenstein-Reichardt Detector (Figure 1A), originally developed to describe behaviour in insects (Hassenstein and Reichardt, 1956), and the Motion Energy Model (Figure 1B), originally developed to explain human perception(Adelson and Bergen, 1985). Both models use nonlinear operators to combine the outputs of several spatial and temporal filters and obtain a direction-sensitive measure of motion strength, known as motion energy, via a final opponent step (Figure 1). This opponent step ensures that they respond only to directional motion, and not to non-directional changes in luminance such as counterphase-modulation or flicker (Qian et al., 1994).

**Figure 1:**
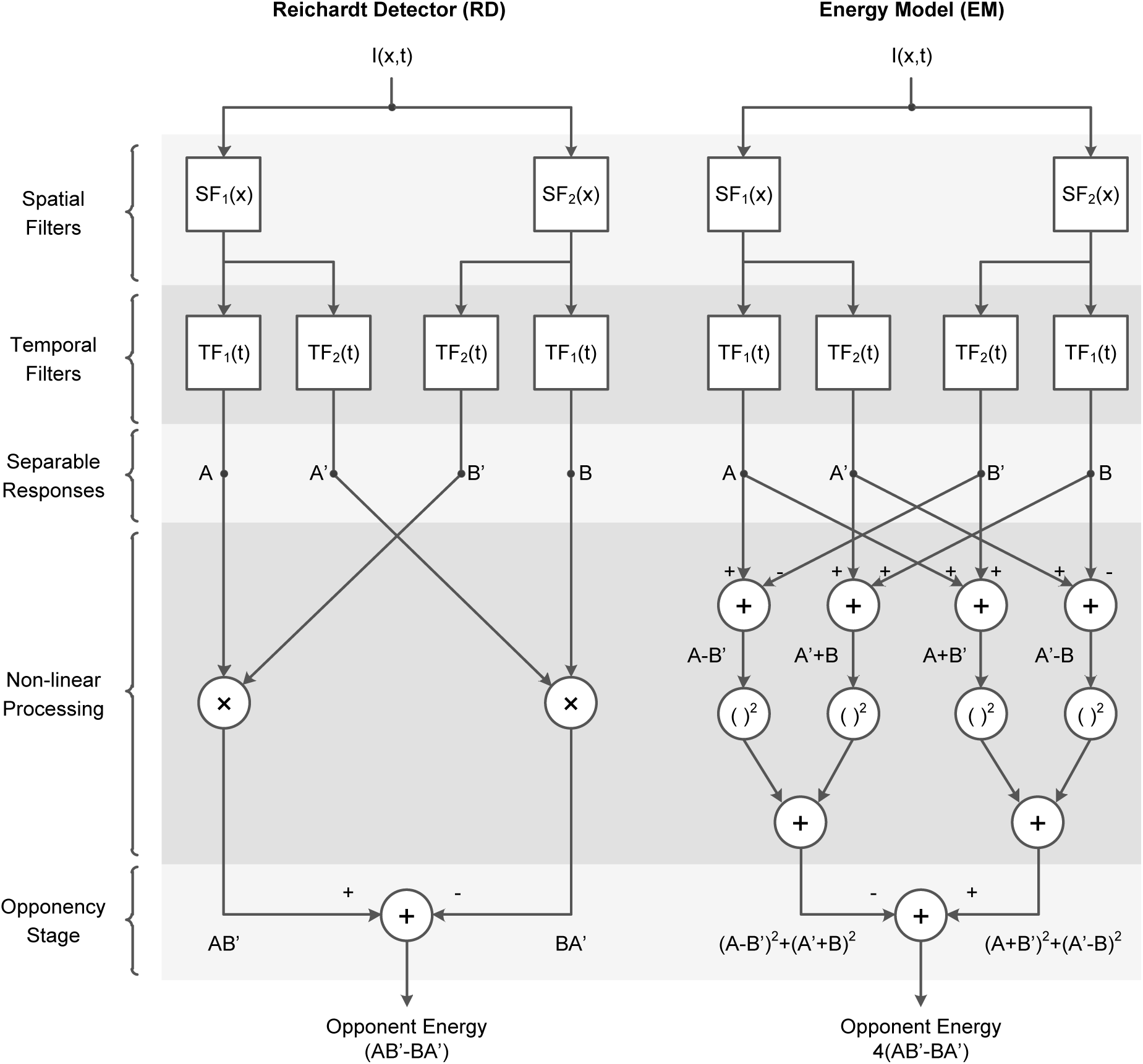
Opponent energy models of motion detection. The Reichardt Detector (RD) and the Energy Model (EM) are two prominent opponent models in the literature of insect and mammalian motion detection. The two models are formally equivalent when the spatial and temporal filters are separable (as shown) and so their outputs and response properties are identical even though their structures are different. Both models use the outputs of several linear spatial and temporal filters (*SF*_1_, *SF*_2_, *TF*_1_ and *TF*_2_) to calculate two opponent terms and then subtract them to obtain a direction-sensitive measure of motion (opponent energy). Nonlinear processing is a fundamental ingredient of calculating motion energy and so both models include nonlinear operators before the opponency stage (multiplication in the RD and squaring in the EM). (Reproduction of Fig. 18 from Adelson and Bergen (1985).)

The two models are traditionally associated with particular early spatiotemporal filters. For the energy model, the filters are often taken to be building blocks for two quadrature pairs of oriented linear responses (Adelson and Bergen, 1985). The attraction of this assumption is that it makes leftward and rightward responses to a simple moving grating constant (despite the temporal modulation of the stimulus) consistent with the existence of directionally-selective complex cells in primary visual cortex (Emerson et al., 1992) and psychophysical evidence for directionally-selective motion detection channels in humans (Levinson and Sekuler, 1975). For the Reichardt detector, the spatial filters are usually assumed to be identical but displaced in space, reflecting the view that their correlates are two neighboring ommatidia in an insect’s eye (Borst, 2014). These filter choices originate from the studies in insects and mammals where the models have their historical roots, but are not intrinsic features of the models themselves.

In fact, although the circuits originally proposed for the motion energy and Reichardt detectors are structurally different (Figure 1), if the same spatiotemporally separable filters are used in each model, the output of the two models is mathematically identical (Adelson and Bergen, 1985; Borst and Helmstaedter, 2015; Lu and Sperling, 1995; Van Santen and Sperling, 1984). Critically, as Figure 1 shows, both models involve opponency, i.e. they compute the difference between motion energy in opposite directions, either explicitly or implicitly. Our discussion and conclusions will therefore apply to both models equally.

### 3.2 Opponency in motion perception

One feature of opponent energy models of motion detection is that the *spatiotemporal filters* and the *detector itself as a unit* may have different spatiotemporal tuning. This can be illustrated by considering the response of an motion energy detector to a single drifting sinusoidal grating with the contrast function:

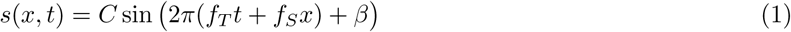

where *x* is the horizontal position of a point in the grating, *t* is time, *f_T_* is temporal frequency, *f_S_* is spatial frequency, *β* is the grating’s phase and *C* is contrast. Following the schematics in Figure 1, both models integrate this stimulus over space and then pass the results through temporal filters to generate the separable time responses:

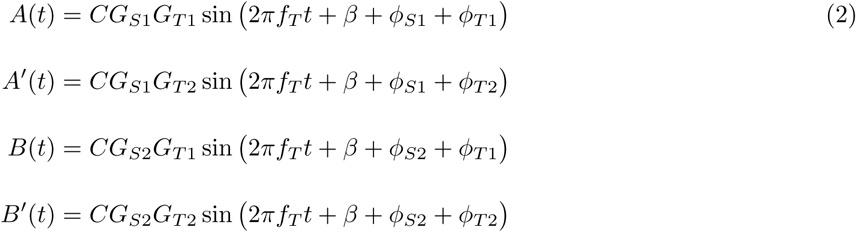

where *G_T_*_1_, *G_T_*_2_, *ϕ_T_*_1_, *ϕ_T_*_2_ are the gains and phase responses of temporal filters at the stimulus temporal frequency *f_T_* and *G_S_*_1_, *G_S_*_2_, *ϕ_S_*_1_, *ϕ_S_*_2_ are likewise for the spatial filters. In the energy model (Figure 1B), these signals are combined into distinct rightward and leftward terms:

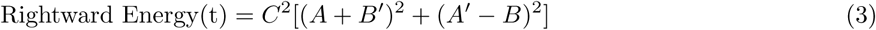

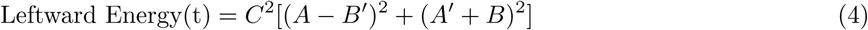

that are subtracted to produce the model output: opponent energy. The Reichardt detector combines the separable responses differently (Figure 1A) but produces the same output (up to a scaling factor of 4):

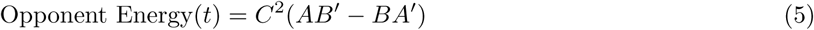

Figure 2 illustrates the Fourier spectrum of rightward, leftward and opponent energy for typical human and insect filters. The red and blue lines in Figure 2AB mark the passband of rightward and leftward energies respectively (Equations 3 and 4). Figure 2A does this for filters designed to model human vision, while Figure 2B is for filters designed to model insect vision; see Methods for details. Figure 2CD shows the opponent energy (defined as rightward minus leftward energy, Equation 5), which is the output of the motion detector.

**Figure 2:**
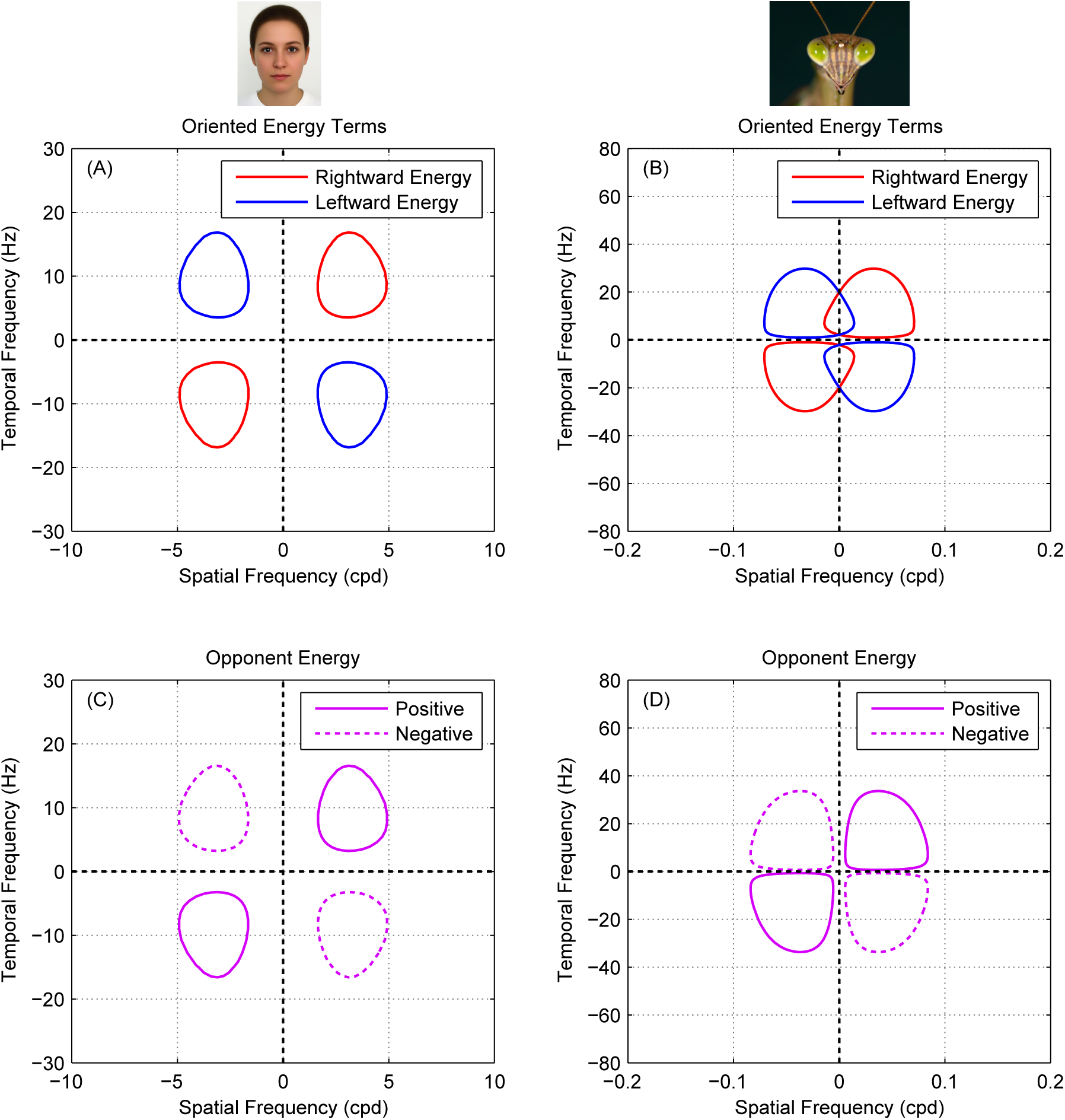
Spatiotemporal filter and opponent energy tuning in an opponent motion model. (**A, B**) Fourier spectra of rightward and leftward energies ((*A* + *B′*)^2^ + (*A′* – *B*)^2^, Equation 3, and (*A* – *B′*)^2^ + (*A′* + *B*)^2^, Equation 4) for example mammalian and insect opponent energy motion detectors (see Methods for details, Equations 15–19). The colored lines in each plot are 0.25 sensitivity contours. The Fourier spectra of leftward and rightward energies are very similar to the model’s filters in each case: spatially bandpass in mammals and low-pass in insects. (**C, D**) Opponent energy, *AB′* – *A′B*, computed as the difference between rightward and leftward energies. In mammals, rightward and leftward responses do not overlap because the spatial filter are band-pass (panel A). In insects, the low-pass spatial filters cause an overlap between rightward and leftward responses (panel B) but this overlap is canceled at the opponency stage making opponent energy insensitive to low spatial frequencies (panel D).

For mammals, early spatiotemporal filters are typically relatively narrow-band, with little response to DC (Anderson and Burr, 1989). The rightward and leftward energies are therefore also bandpass and clearly separated in Fourier space (Figure 2A), very similar to those of the input filters. The regions of Fourier space where the opponent energy is positive (bounded by solid contours in Figure 2C) are simply the same regions where there is rightward energy (bounded by red in Figure 2A), and similarly for negative/leftward (dotted in C, blue in A). Thus, there are no frequencies that elicit a strong response from the individual filters and not from the opponent model as a whole.

For insects, the situation is different. The two spatial inputs to a Reichardt detector are usually taken to be a pair of adjacent ommatidia (Buchner, 1976; Pick and Buchner, 1979), so the spatial filter is simply the angular sensitivity function of an ommatidium, which is lowpass, roughly Gaussian (Rossel, 1979; Van Santen and Sperling, 1984). Accordingly, as shown in Figure 2B, insects have substantial leftward and rightward energy responses at zero spatial frequency. Crucially, these are canceled out in the opponency step, meaning that the Reichardt detector as a whole does not respond to whole-field changes in brightness to which individual photoreceptors do respond. Thus the opponent energy terms are bandpass (Figure 2D). This means that, for insects, the spatiotemporal filters and the model itself as a unit may have different spatiotemporal tuning.

Mathematically, after substituting for the filter outputs in Equation 5 and simplifying, the output of the motion detector can be expressed as

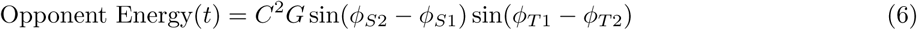

where *G* is the product of the filter gains, *G* = *G_S_*_1_ *G_S_*_2_ *G_T_*_1_ *G_T_*_2_, and the *ϕ* are the phases of the filter responses, defined above in Equation 2. Since *G* is spacetime separable, it is not direction-selective. The direction-selectivity is created by the phase-difference terms. Since the filters are real, the filter phase is an odd function of frequency. This means that the energy is positive in the first and third Fourier quadrants and negative in the second and fourth, as shown in Figure 2CD.

The important point for our purposes is that the frequency tuning of the motion detector as a whole reflects both that of the filter gains *G*, *and* that of the phase-difference terms. For the Reichardt detector, the phase-difference terms make the motion detector spatially bandpass even though its spatial filters are lowpass. In the Reichardt detector, the spatial filters are identical but offset in position by a distance Δ*x*, so the spatial phase-difference term in Equation 6 is sin(2*πf_S_*Δ*x*). This term removes the response to the lowest frequencies, as we saw in Figure 2D.

In the energy model, the spatial filters are usually taken to be bandpass functions like Gabors or derivatives of Gaussians, differing in their phase but not position. For such functions, the phase difference is independent of frequency, so the phase-difference terms in Equation 6 just contribute an overall scaling and the frequency tuning of the motion detector is determined solely by the filter gains G. This remains approximately true even for filters which differ in position as well as phase, provided they are bandpass. We shall show that this difference in the bandwidth of their spatial filters means that the energy model and Reichardt detector are affected very differently by motion noise, despite the fact that the model architecture is mathematically identical.

### 3.3 Response to a general stimulus

We now work through what happens when noise is added to a motion signal. We consider the response of an opponent model to an arbitrary stimulus composed of a sum of *N* drifting gratings:

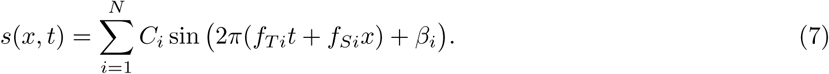

Since the filters in an energy opponent motion detector are linear, the separable responses *A, A′, B* and *B′* can be expressed as a sum of the independent responses to the components present in a stimulus. The model’s overall response to the compound grating in Equation 7 can therefore be written as:

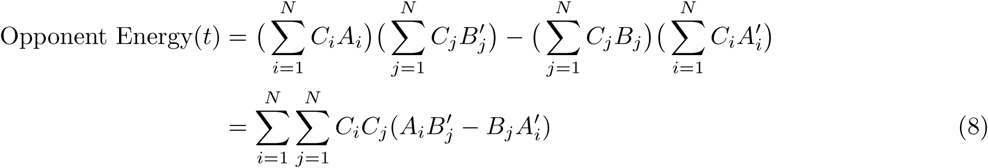

where the subscripts denote responses to the components. To simplify, we extract the terms where *i* = *j* and re-write the expression as:

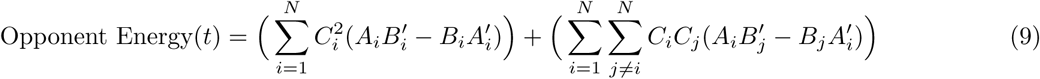

The response in Equation 9 consists of two parts. Terms within the first sum operator (the *independent terms*) are simply the summed responses to grating components when presented each on its own (Equation 5). Obviously, frequencies which do not elicit a response when presented in isolation do not contribute to this term. The remaining terms within the second sum operator represent crosstalk or *interactions* between component pairs at different spatial and/or temporal frequencies. These show more subtle behaviour.

Interactions differ from independent terms in a number of ways. First, if two components have different temporal frequencies then their interaction is a sinusoidal function of time, so has no net contribution to the response when integrated over time (Van Santen and Sperling, 1984, 1985). When two components *i* and *j* have the same temporal frequency, however, their interaction results in the DC response:

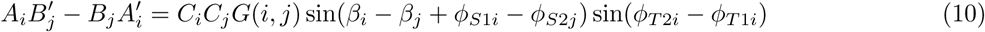

where *G*(*i, j*) = *G_S_*_1*i*_*G_S_*_2*j*_*G_T_*_1*i*_*G_T_*_2*i*_, the product of the filter gains at the spatial and temporal frequencies in question. *β_i_*, *β_j_* are the phases of the stimulus components (Equation 7), *ϕ_S_*_1*i*_, *ϕ_S_*_2*j*_ are the phases of the two spatial filters at the relevant frequencies, *f_Si_* and *f_S_j*, and *ϕ_T_*_1*i*_, *ϕ_T_*_2*i*_ are the phases of the two temporal filters at the temporal frequency *f_Ti_*. This response has a similar form to Equation 6 but differs in an important way: its spatial phase-difference term depends on the spatial filter phase responses *to different stimulus components*. Suppose there is a spatial frequency *f_Sj_* for which both spatial filters have substantial gains *G_S_*_1*j*_, *G_S_*_2*j*_ and equal phases: *ϕ_S_*_1*j*_ = *ϕ_S_*_2*j*_. Due to opponency, this component will not elicit any response when presented in isolation, because of the term sin(*ϕ_S_*_1*j*_ – *ϕ_S_*_2*j*_) in Equation 6; it will appear invisible to the detector. Yet its interaction with a visible component *f_Si_* will nevertheless add a constant offset to the model’s output, provided only that sin(*β_i_* – *β_j_*+ *ϕ_S_*_1*i*_ – *ϕ_S_*_2*j*_) ≠ 0. This means that invisible noise at *f_Sj_* can mask a signal at *f_Si_*.

### 3.4 Early spatial filtering in insect vs mammalian motion detection

Does this effect actually occur in biological motion detectors? In mammals, it seems the answer is no. There, the spatial filters are bandpass functions like narrow-band Gabors or derivatives of Gaussians, which have roughly constant phase for all frequencies of a given sign. The two spatial filters are generally modelled as having the same position but different phase, which means that there are no components for which *ϕ_S_*_1*j*_ = *ϕ_S_*_2*j*_. If the filters had different positions as well as phases, such components could exist, but this would imply some strange properties of the motion detector (tuning to different directions for different frequency components) which have not been reported. For realistic mammalian filters, therefore, it is not possible for components to be invisible when presented in isolation and yet to affect the response to visible components.

However, for insect motion detectors, the spatial filters are believed to resemble Gaussians with a spatial offset Δ*x*. For a component with spatial frequency *f_Sj_*, the phase difference between the two filters is 2*π*Δ*xf_Sj_*. As the spatial frequency tends to zero, so does the phase difference and thus the response of the opponent energy motion detector (Equation 6). The opponent energy detector as a whole is therefore bandpass in its spatial frequency tuning, as has been confirmed many times for insects (Borst, 2014; Dvorak et al., 1980; Nityananda et al., 2015; OCarroll et al., 1997; OCarroll, DC and Bidwell, NJ and Laughlin, SB and Warrant, EJ, 1996). Yet since the Gaussian filters are low-pass, the gain *G_S_*2_*j*_ remains high. This means that there can be a large interaction term between this frequency and visible frequencies *f_Si_* (Equation 10).

This analysis suggests that the interaction terms produced by the nonlinearity of the motion energy model can indeed be safely ignored for mammals, so long as the relevant spatial filters are bandpass. However, we predict that in insects, motion signals can be masked by invisible noise. This effect has not so far been demonstrated.

### 3.5 Mammalian motion detection is not affected by invisible noise

The spatiotemporal frequency tuning of motion detectors is often estimated psychophysically by measuring their responses to masked gratings. In these experiments, detecting a coherently-moving grating (the signal) is made more difficult by superimposing one or more gratings with different spatial/temporal frequencies but no coherent motion (the noise). The relative increase in detection threshold as a function of noise frequency (i.e. the masking function) is taken as the spatiotemporal sensitivity of the individual detector (Anderson and Burr, 1989). This technique is important because it enables the tuning of a single channel to be inferred, even though many channels contribute to the spatiotemporal sensitivity of the whole organism.

Figure 3 reproduces data from (Anderson and Burr, 1985) showing such an experiment in humans. The reduction in sensitivity is greatest when noise is at the same spatial frequency as the signal. As noise moves away from the signal frequency, either higher or lower, it has progressively less effect. In this way, Anderson and Burr (1985) deduced that human motion channels are bandpass with a bandwidth of 1 to 3 octaves.

**Figure 3:**
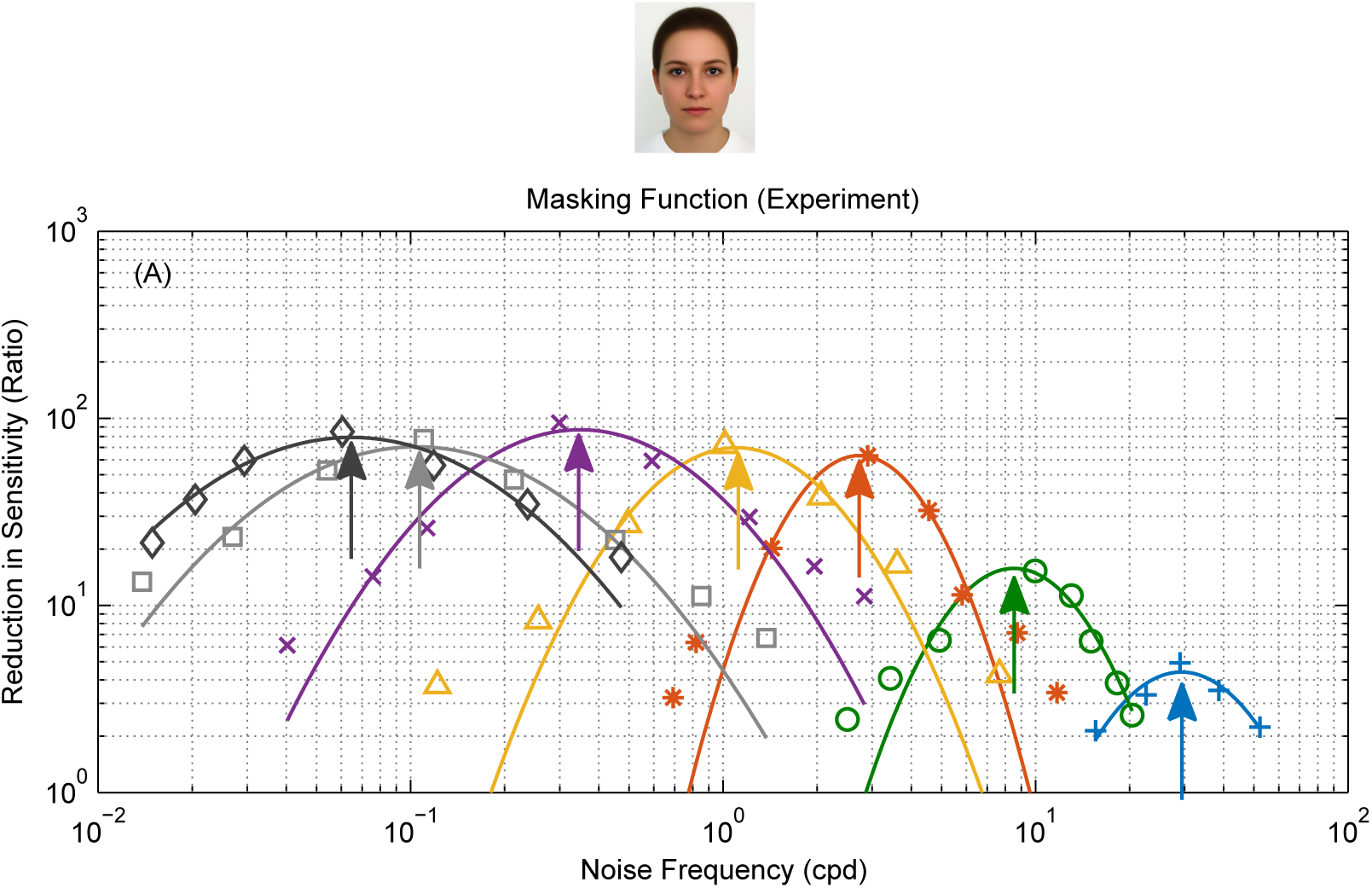
Effect of noise on mammalian motion detectors. Measurements showing the effect of noise on motion detection sensitivity in humans (reproduction of Fig. 1b from Anderson and Burr (1985)). The colored plots show responses to different signal frequencies (marked by arrows). Noise is most effective at masking the signal when its frequency is the same and less effective as its frequency changes in either direction.

We model this by assuming that motion is detected when the output of an motion detector exceeds a threshold (see Methods for details). Because noise carries no motion signal, it has no effect on the mean detector output, but it increases its variability and hence decreases the proportion of above-threshold responses. This leads to a decrease in response rate and consequently threshold elevation. The factor by which threshold is elevated for noise at a given frequency forms the masking function, whose shape reflects the variability of the motion detector output.

Figure 4 shows the results of this simulation. Figure 4A shows the spatial sensitivity function of an energy model motion detector, i.e. its response to single drifting gratings as a function of their spatial frequency. This is also the detector’s mean response in the presence of noise. However, noise elevates the variability of the response, as shown in Figure 4C. Accordingly, the signal contrast needed for the model to reliably detect motion is increased, and we obtain the masking function shown in Figure 4D. As Anderson and Burr (1985) assumed, this accurately reflects the sensitivity of the underlying mechanisms (cf. Figure 4D and Figure 4A). In particular, the bandpass filter tuning gives bandpass masking.

**Figure 4:**
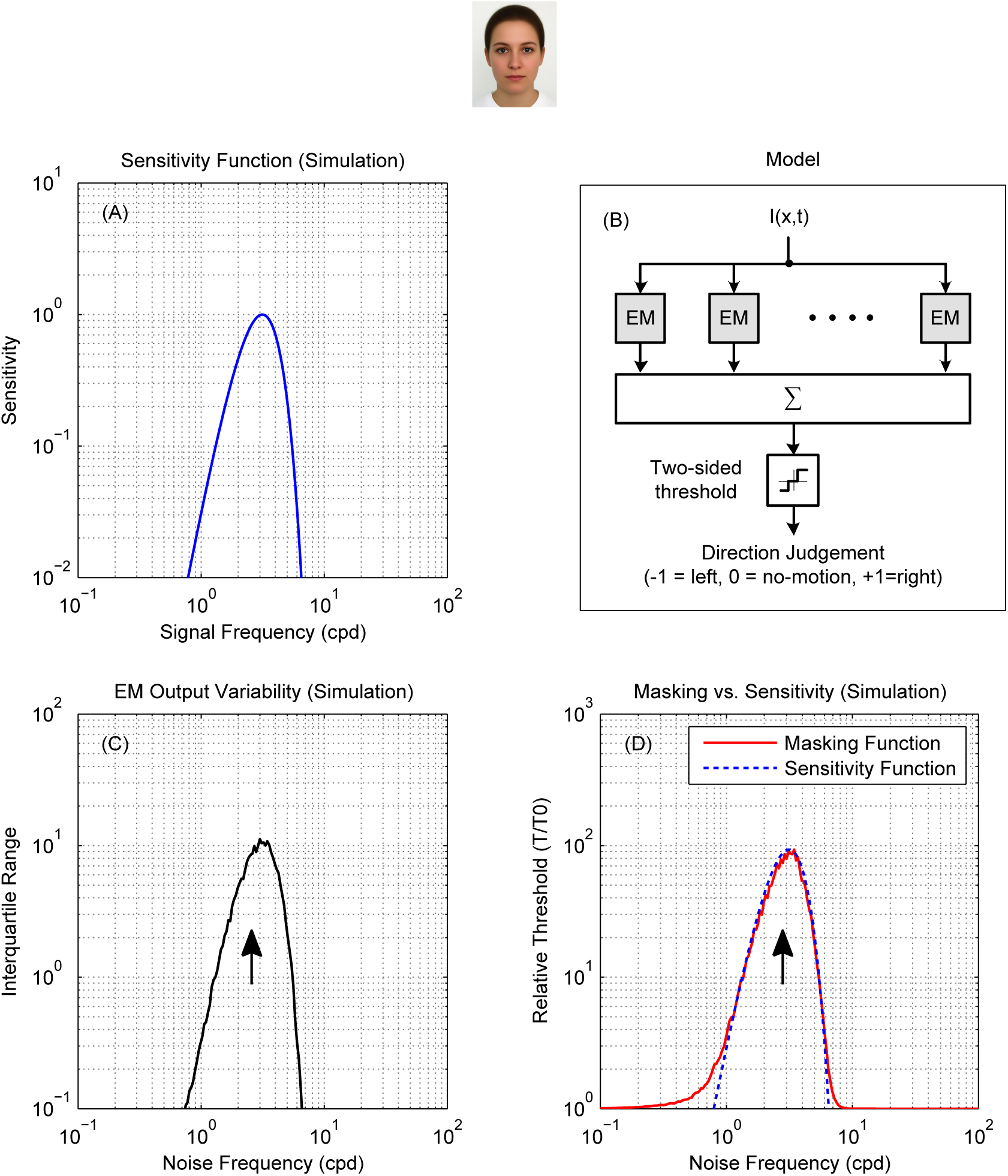
For mammalian bandpass filters, the masking function reflects sensitivity. (**A**) (**B**) Direction discrimination model based on an array of opponent models with the spatial tuning plotted in panel A. Opponent model outputs are pooled and passed through a two-sided threshold of value *T* to produce a ternary judgment of motion direction per stimulus presentation. (**C**) The variability of opponent model outputs across 500 simulated presentations (per noise frequency point) of a noisy stimulus consisting of a signal grating of 3 cpd and temporally-broadband noise. Signal frequency is marked on the plot with an arrow. Signal and noise had 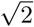 and 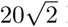 RMS contrast respectively. Adding noise did change the mean of opponent output but had a significant effect on its spread. Output variance was highest when noise frequency was 3 cpd and lower as noise frequency changed in either direction, closely resembling the shape of the opponent model’s sensitivity function. (**D**) The masking function (red) was calculated based on these simulated results as *T*(*f_n_*)/*T*_0_ where *T*(*f_n_*) is the threshold corresponding to a 90% detection rate at each noise frequency and *T*_0_ is the detection threshold of an unmasked grating. The sensitivity function from (A) is reproduced, scaled, for comparison (blue dotted line). The masking function is a good approximation to the sensitivity.

Thus in mammals, the masking function can be used to infer (approximately) the spatiotemporal sensitivity of motion detection channels. This works because the initial filters are spatiotemporally bandpass (Anderson and Burr, 1985, 1989; Burr et al., 1986a,b).

### 3.6 Insect motion detection is affected by invisible noise

As we have seen, the response of insect motion detectors to masked grating stimuli is is expected to be qualitatively different. The lowpass tuning of the early spatial filters in models of insect motion detection predicts that low-frequency components which elicit no response when presented on their own will still influence the detector’s response to other frequencies.

This means that for insects, we predict differences between their motion masking and sensitivity functions. Specifically, when the mask contains components with the same temporal frequency as the signal, we expect the masking effect of noise to extend to spatial frequencies much lower than the sensitivity band of an insect motion detector. In this section, we present the results of experiments in which we tested this prediction.

Figure 5A shows the mantis optomotor response rates we measured in our experiment as a function of noise frequency. For insects, trials are slow, so we did not attempt to measure contrast thresholds for each combination of signal and noise. Rather, we measured response rates at a single signal contrast, and used the drop in response rate to assess the effect of noise. To facilitate comparison with the corresponding plot in mammals (Figure 3A), we plot the *masking rate*, defined as *M*(*f_n_*) = (*R*_0_ – *R*(*f_n_*))/*R*_0_ where *R*(*f_n_*) is the optomotor response rate at a given noise frequency and *R*_0_ is the baseline response rate (measured without adding noise).

**Figure 5:**
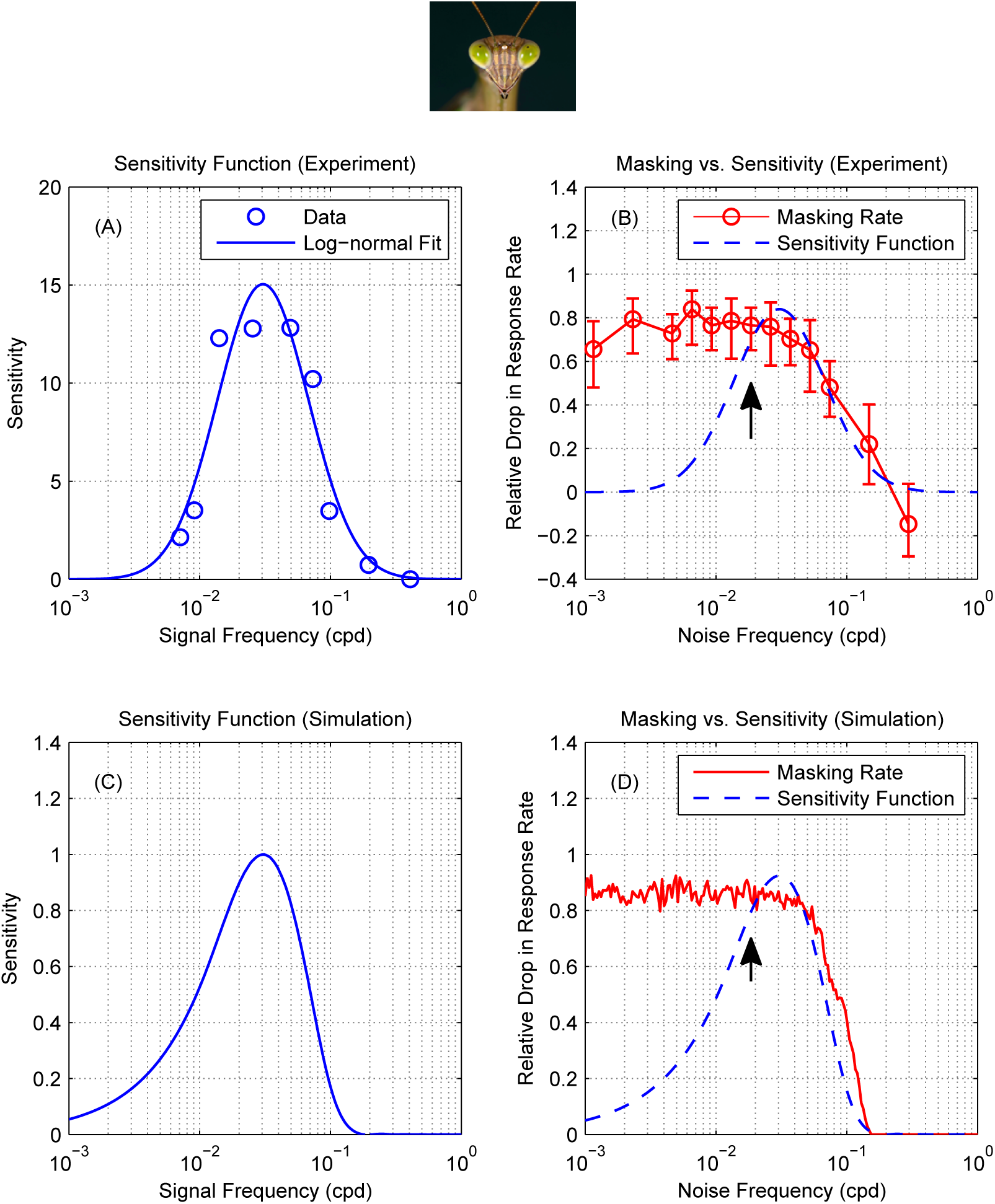
For mantis motion detection, masking function does not reflect sensitivity. (**A**) The spatial sensitivity of mantis motion detectors at 8 Hz, measured using the same experimental paradigm, showing bandpass sensitivity in the range 0.01 to 0.1 cpd (reproduction of Fig. 3a in Nityananda et al. (2015)). (**B**) Measurements showing the effect of noise on the detection of a moving grating in the praying mantis. Circles are masking rate *M* defined as *M* = (*R*_0_ – *R*)/*R*_0_ where *R* is the response rate (proportion of trials in which mantids responded optomotorally in the same direction as the signal grating) and *R_0_* is the baseline (no-noise) response rate. Error bars are 95% confidence intervals calculated using simple binomial statistics. Signal frequency (0.0185 cpd) is marked on the plot with an arrow. The response rate measured at 0.03 cpd was slightly below baseline and so the calculated masking rate was negative. (**C**) Normalized sensitivity function of a motion energy model tuned to 0.03 cpd (18). (**D**) Simulated masking function (red) with the simulated sensitivity function reproduced for comparison (blue dotted line, scaled to same peak). Masking and sensitivity functions in the mantis are qualitatively different: noise below the lower end of the sensitivity function (˜ 0.01 cpd) continues to mask the signal.

As in mammals, adding noise to the stimulus causes a drop in response rate (corresponding to an increase in the masking rate). Unlike mammals, however, the impact of masking on the mantis is not predicted by its motion sensitivity function. For example, injecting noise near the peak spatial sensitivity (0.03 cpd, Nityananda et al. (2015)) unsurprisingly causes severe masking; the masking rate is 80%. In the absence of noise, with a signal of contrast 0.125 at 0.0185 cpd, insects moved in the direction of the signal on *R*_0_ = 60% of trials; after adding noise with contrast 0.198 at 0.03 cpd, this dropped to *R*(*f_n_*) = 12%. Also unsurprisingly, injecting noise at frequencies much higher than the peak has little effect. For example, noise at 0.3cpd produces masking which is not significantly different from zero; this is expected given that Nityananda et al. (2015) found sensitivity at 0.3 cpd was near zero (their Fig 3b).

But counter-intuitively, noise injected at frequencies much lower than the peak continues to produce strong masking. For example, Nityananda et al. (2015) found that sensitivity at 0.007 cpd, the lowest frequency they tested, was 15% of the peak value. Normally, we would expect the effect of noise to be reduced correspondingly. However, we find the masking rate at 0.0025 cpd is still 80%, just as severe as noise injected at the peak.

We tested the effect of noise on three further signal frequencies (Figure 6). The amount of masking depends on the signal frequency. Since we always presented the signal grating at the same contrast, the effective strength of the signal depends on the sensitivity at the signal frequency. Accordingly, noise has least effect on signals at 0.037 cpd (maximum masking rate 50%, Figure 6A), and most effect at the lowest and highest signal frequencies (maximum masking rate 80% for 0.0185 cpd, Figure 5A, and 95% for 0.177 cpd, Figure 6C). As the noise frequency increases much beyond the signal frequency, it produces progressively less masking. The precise frequency at which the high frequency fall-off occurs depends on the signal frequency. This may reflect the contribution of motion detectors with different selectivity for spatial frequencies

**Figure 6:**
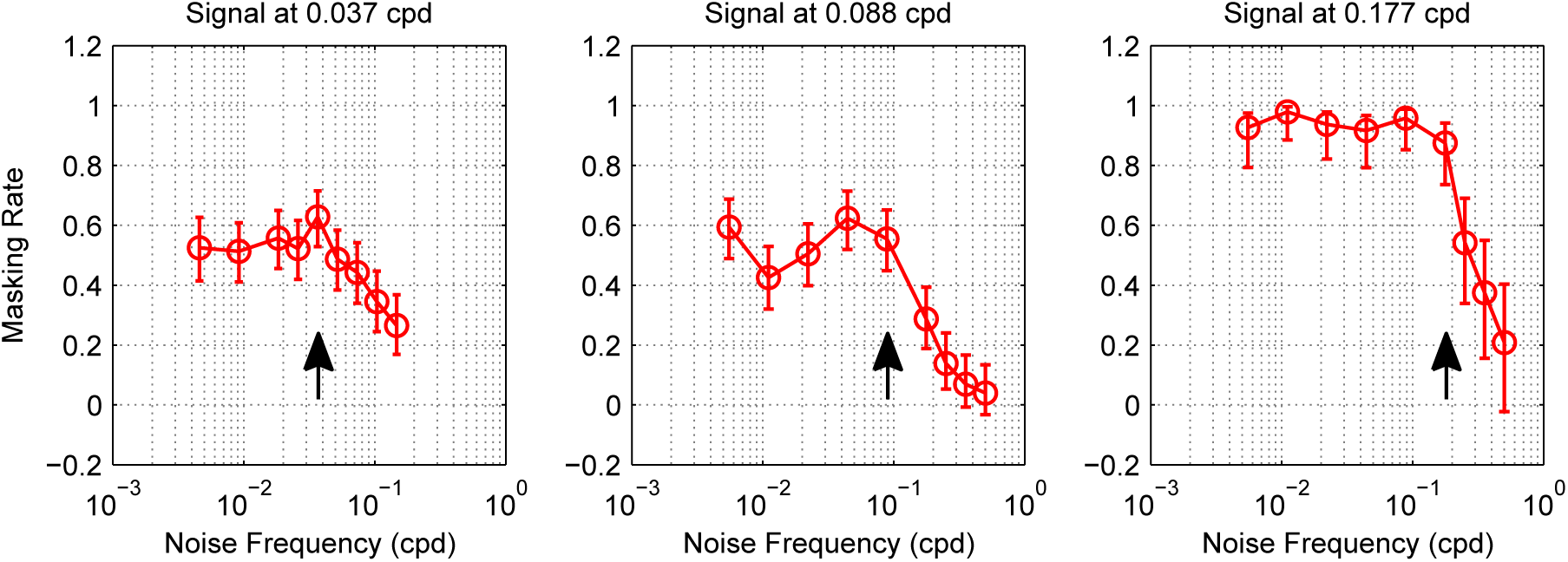
Mantis masking rate measurements at different signal frequencies. Measurements of masking rate versus noise frequency in the mantis (for the signal frequencies 0.037, 0.088 and 0.177 cpd) showing the same masking trends as Figure 5A (signal frequency 0.0185 cpd): noise continues to mask the signal significantly even if its frequency is below the spatial sensitivity passband of mantis motion detectors (~ 0.01 to 0.1 cpd). Circles are masking rate M defined as *M* = (*R*_0_ – *R*)/*R*_0_ where *R* is the response rate (proportion of trials in which mantids responded optomotorally in the same direction as the signal grating) and *R*_0_ is the baseline (no-noise) response rate. Error bars are 95% confidence intervals calculated using simple binomial statistics. Signal frequency is marked on each plot by an arrow.

Critically, in every case, we found the same dependence on noise frequency: noise injected at frequencies at or below the signal frequency produced essentially the same amount of masking, regardless of the precise frequency it was injected at. That is, masking is low-pass, even though the insects’ motion sensitivity is bandpass. This is the signature interaction effect we predicted we would find in insects, due to their lowpass early spatial filtering combined with the nonlinearity of the standard models of motion detection.

## 4 Discussion

We show that the standard model of motion detection produces nonlinear interactions between the spatial components of moving stimuli. Stimuli that elicit no response can nonetheless have a powerful masking effect, if the filters that precede motion detection are spatially lowpass. We show that this sort of mask does effectively disrupt the optomotor response of the praying mantis. This is very different from the effects of masking noise in humans, but our analysis suggests that this reflects the same motion computation in both species, computed after different initial filters are applied. This highlights the fact that simple nonlinearities can have complex effects. In human studies, it is commonly assumed that nonlinear interactions take place only within the sensitivity band of a given channel within a system (Anderson and Burr, 1989; Daugman, 1984), where a “channel” is a pool of neurons with similar tuning (Blakemore and Campbell, 1969; Campbell and Robson, 1968; De Valois and Tootell, 1983; Graham and Nachmias, 1971; Sachs et al., 1971). This turns out to be a good approximation only if the sensitivity band is set by the inputs to the channel, rather than by subsequent nonlinearities. This is true for humans, but not in insects.

Here, we have analysed the standard model of motion detection. This is mathematically equivalent to both the Reichardt Detector and to the Motion Energy Model, the standard accounts of motion detection in insects and mammals respectively (Anderson and Burr, 1989; Hassenstein and Reichardt, 1956). The two accounts have different circuitry but are mathematically equivalent when the same filters are used as inputs (Adelson and Bergen, 1985; Borst and Helmstaedter, 2015; Lu and Sperling, 1995; Van Santen and Sperling, 1984). We derived equations 9 and 10 showing how such motion detectors can be affected by frequency components outside their sensitivity band (Chen et al., 1993). In these models, interaction terms with different temporal frequencies average to zero over time, producing “pseudo-linearity” (Van Santen and Sperling, 1984). Crucially, however, we show that cross-frequency interactions can survive opponency and time-averaging. When low spatial frequencies are transmitted by early spatiotemporal filters, even if they are normally cancelled subsequently by the opponency step, these “invisible” components can affect the response to other, visible signals.

For mammalian motion sensors, this effect may be a mathematical curiosity. Motion sensors are built at a relatively late stage, following early neural filtering (Morgan, 1992) which is spatially bandpass for both spatial and temporal frequency, even as early as the output of the retina. The opponency in models of mammalian motion detection sharpens direction selectivity, but has little effect on spatial frequency tuning. In contrast, current models of insect motion sensors postulate that they are constructed at a much earlier stage, directly from individual ommatidia. The filters are spatially-lowpass, reflecting largely optical, rather than neural, factors (Rossel, 1979; Snyder et al., 1977). Our analysis predicted that this would make insect motion detectors subject to interference from “invisible” low-frequency noise. We have confirmed this behaviourally in an insect model, the praying mantis.

Given the differences between humans and mantises, it is remarkable that the experimental data in both species is so well described by a model of exactly the same structure (Figure 4, Figure 5). This model employs a simple decision rule in which motion is perceived when the average activity of a group of motion detectors exceeds a threshold. The only difference is the early spatiotemporal filters used for each species: spatially bandpass for mammals and spatially lowpass for insects. In both cases the masking function reflects this early spatial filtering. For mammals, this is the same as the spatiotemporal sensitivity of the whole organism, but for insects it is not. Thus, the same circuitry results in very different behaviour.

Although motion perception presumably evolved independently in insects and mammals, the underlying circuits may be much older. The circuit relies on two very common operations: an output nonlinearity and a subtraction. These are both very common operations, so similar circuits are likely to be widespread in nervous systems. These common operations can lead to very different behaviour, given only slight differences in the inputs. It seems likely that other behavioural differences may be explained in equally simple ways.

## 5 Materials & Methods

We used a masking paradigm to test visual motion detection in the praying mantis. In the context of motion detection, a “signal” is an image that moves smoothly in a given direction, to “detect the signal” is to report the direction of motion and “noise” is a sequence of images with no consistent motion. mantises were placed in front of a CRT screen and viewed full screen gratings drifting either leftward or rightward in each trial. In a subset of trials, the moving grating elicited the optomotor response, a postural stabilization mechanism that causes mantises to lean in the direction of a moving large-field stimulus. An observer coded the direction of the elicited optomotor response in each trial (if any) and these responses were later used to calculate motion detection probability as the proportion of trials in which mantises leaned in the same direction as the stimulus. Videos of mantises responding optomotorally to a moving grating using same experimental paradigm are available on (http://www.edge-cdn.net/video_839277?playerskin=37016) and (http://www.edge-cdn.net/video_839281?playerskin=37016) (supplementary material to Nityananda et al. (2015)).

### 5.1 Insects

The insects used in experiments were 11 individuals (6 males and 5 females) of the species *Sphodromantis lineola*. Each insect was stored in a plastic box of dimensions 17 × 17 × 19 cm with a porous lid for ventilation and fed a live cricket twice per week. The boxes were kept at a temperature of 25° C and were cleaned and misted with water twice per week.

### 5.2 Experimental Setup

The setup consisted of a CRT monitor (HP P1130, gamma corrected with a Minolta LS-100 photometer) and a 5 × 5 cm Perspex base onto which mantises were placed hanging upside down facing the (horizontal and vertical) middle point of the screen at a distance of 7 cm. The Perspex base was held in place by a clamp attached to a retort stand and a web camera (Kinobo USB B3 HD Webcam) was placed underneath providing a view of the mantis but not the screen. The monitor, Perspex base and camera were all placed inside a wooden enclosure to isolate the mantis from distractions and maintain consistent dark ambient lighting during experiments.

The screen had physical dimensions of 40.4 × 30.2 cm and pixel dimensions of 1600 × 1200 pixels. At the viewing distance of the mantis the horizontal extent of the monitor subtended a visual angle of 142°. The mean luminance of the stimuli was 51.4 cd/m^2^ and its refresh rate was 85 Hz.

The monitor was connected to a Dell OptiPlex 9010 (Dell, US) computer with an Nvidia Quadro K600 graphics card and running Microsoft Windows 7. All experiments were administered by a Matlab 2012b (Mathworks, Inc., Massachusetts, US) script which was initiated at the beginning of each experiment and subsequently controlled the presentation of stimuli and the storage of keyed-in observer responses. The web camera was connected and viewed by the observer on another computer to reduce processing load on the rendering computer’s graphics card and minimize the chance of frame drops. Stimuli were rendered using Psychophysics Toolbox Version 3 (PTB-3) (Brainard, 1997; Kleiner et al., 2007; Pelli, 1997).

### 5.3 Experimental Procedure

Each experiment consisted of a number of trials in which an individual mantis was presented with moving gratings of signal and noise components. An experimenter observed the mantis through the camera underneath and coded its response as “moved left”, “moved right” or “did not move”. The camera did not show the screen and the experimenter was blind to the stimulus. There were equal repeats of left-moving and right-moving gratings of each condition in all experiments. Trials were randomly interleaved by the computer. In between trials a special “alignment stimulus” was presented and used to steer the mantis back to its initial body and head posture as closely as possible. The alignment stimulus consisted of a chequer-like pattern which could be moved in either horizontal direction via the keyboard and served to re-align the mantis by triggering the optomotor response.

### 5.4 Visual Stimulus

The stimulus consisted of superimposed “signal” and “noise” vertical sinusoidal gratings. The signal grating had one of the spatial frequencies 0.0185, 0.0376, 0.0885 or 0.177 cpd, a temporal frequency of 8 Hz and was drifting coherently to either left or right in each trial. Signal temporal frequency was chosen to maximize the optomotor response rate based on the mantis contrast sensitivity function (Nityananda et al., 2015). Noise had a spatial frequency in the range 0.0012 to 0.5 cpd and its phase was randomly updated on each frame to make it temporally broadband (with a Nyquist frequency of 42.5 Hz) without net coherent motion in any direction. Each presentation lasted for 5 seconds.

Since mantises were placed very close to the screen (7 cm away), any gratings that are uniform in cycles/px would have appeared significantly distorted in cycles/deg (Anderson and Burr, 1985). To correct for this we applied a nonlinear horizontal transformation so that grating periods subtend the same visual angle irrespective of their position on the screen. This was achieved by calculating the visual degree corresponding to each screen pixel using the function:

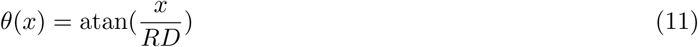

where *x* is the horizontal pixel position relative to the center of the screen, *θ*(*x*) is its visual angle, *R* is the horizontal screen resolution in pixels/cm and *D* is the viewing distance. To an observer standing more than *D* cm away from the screen, a grating rendered with this transformation looked more compressed at the center of the screen compared to the periphery. At *D* cm away from the screen, however, grating periods in all viewing directions subtended the same visual angle and the stimulus appeared uniform (in degrees) as if rendered on a cylindrical drum. This correction only works perfectly if the mantis head is in exactly the intended position at the start of each trial and is most critical at the edges of the screen. As an additional precaution against spatial distortion or any stimulus artifacts caused by oblique viewing we restricted all gratings to the central 85° of the visual field by multiplying the stimulus luminance levels *L*(*x,y,t*) with the following Butterworth window:

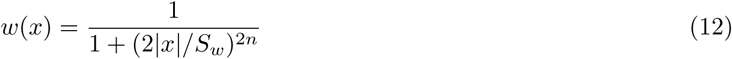

Where *x* is the horizontal pixel position relative the middle of the screen, *S_w_* is the window’s Full Width at Half Maximum (FWHM), chosen as 512 pixels in our experiment (subtending a visual angle of 85° at the viewing distance of the mantis), and n is the window order (chosen as 10). This restriction minimized any spread in spatial frequency at the mantis retina due to imperfections in our correction formula described by Equation 11. We have previously shown that the mantis optomotor response is largely driven by the central visual field, such that a stimulus covering the central 85° should elicit around 84% of the response which would have been elicited by a stimulus covering the entire visual field (Nityananda et al., 2017).

With the above manipulations the presented stimulus was:

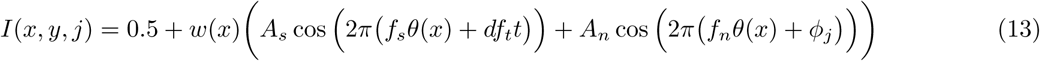

where *x, y* are the horizontal and vertical positions of a screen pixel, *k* is frame number, *I* is pixel luminance, in normalized units where 0 and 1 are the screen’s minimum and maximum luminance levels (0.161 and 103 cd/m^2^ respectively), *A_s_* is signal Michelson contrast (0.125), *A_n_* is noise contrast (0.198), *f_s_* is signal spatial frequency (0.0185, 0.0376, 0.0885 or 0.177 cpd), *f_n_* is noise frequency (varied across trials), *f_t_* is signal temporal frequency (8 Hz), *d* indicates motion direction (either 1 or −1 on each trial), *ϕ_j_* is chosen randomly from a uniform distribution between 0 and 1 on each frame, t is time in seconds (given by *t* = *j*/85), and θ(*x*) is the pixel visual angle according to Equation 11. Still frames, space-time plots and spatiotemporal Fourier amplitude spectra of the stimulus are shown in Figure 7.

**Figure 7:**
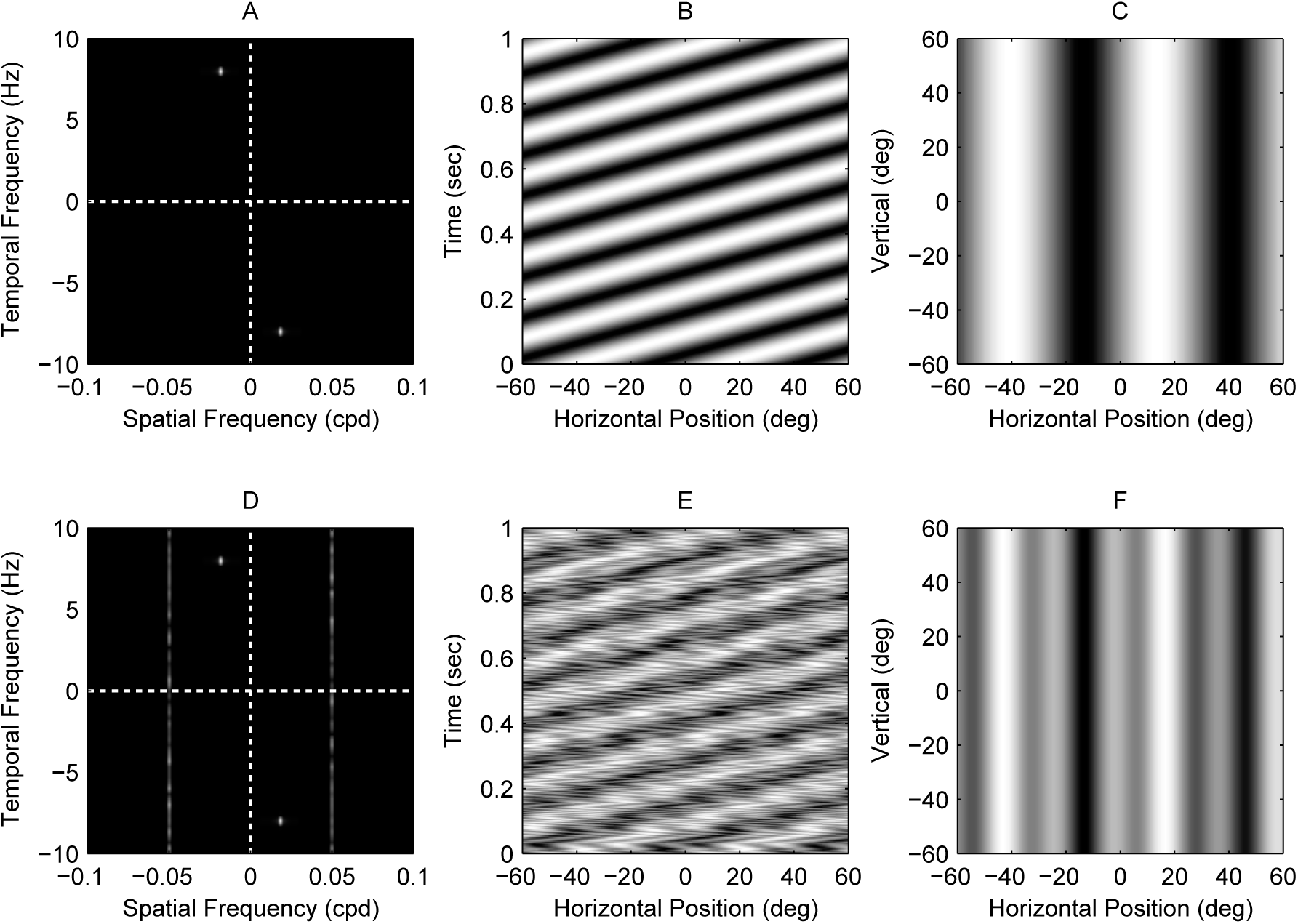
Masked grating visual stimuli used in the experiment. (**A, D**) Spatiotemporal Fourier spectra, (**B, E**) space-time plots and (**C, F**) still frames of the visual stimulus in two conditions of the experiment. Panels A,B,C represent a no-noise condition: the stimulus is a moving grating at 0.0185 cpd and 8 Hz with no added noise. Panels D,E,F represent a masked condition: the stimulus consists of the same signal grating but with non-coherent temporally-broadband noise added at 0.05 cpd. There were in total 44 conditions in the experiment (4 unmasked and 40 masked gratings). Noise was always temporally broadband and its spatial frequency varied across conditions (in the range 0.0012 to 0.5 cpd).

### 5.5 Modeling

Figures 4 and 5 contain simulation results from the model shown in Figure 4 (Panel B). The model consists of 10 opponent energy models (based on the schematics shown in Figure 1 panel B), placed at different positions on a virtual retina, a linear sum and a two-sided threshold of the form:

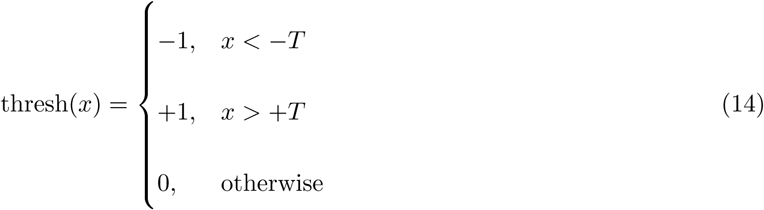

The model was simulated numerically in Matlab. The spatial resolution of simulations was 0.01 deg, time step was 1/85 seconds and each simulated presentation was 1 second long.

The spatial and temporal sensitivity of energy model filters were adjusted to approximate the sensitivities of insects and mammals in different simulations. For mammals (Figure 2AC and Figure 4), spatial filters were second and third derivatives of Gaussians (*σ* = 0.08°) and temporal filters were

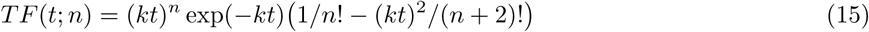

where *n* = 3 for *TF*_1_, *n* = 5 for *TF*_2_ and *k* = 105 for both filters. These filter functions and parameters were taken from the published literature on human motion perception and spatiotemporal tuning (Adelson and Bergen, 1985; Robson, 1966). For insects (Figure 2BD and Figure 5), we used Gaussian spatial filters and first-order low/high pass temporal filters:

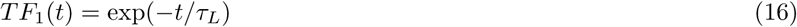

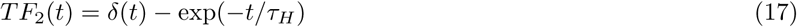

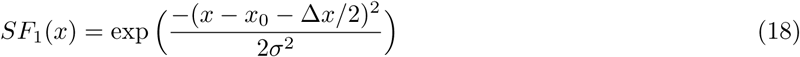

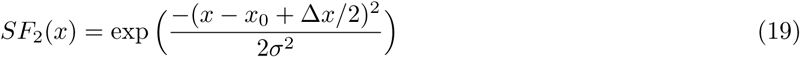

where *τ_L_* = 13 ms, *τ_H_* = 40 ms, Δ*x* = 4°, *σ* = 2.56°. Insect filter functions and parameters were again taken from the published literature (Borst, 2014) (Rossel, 1979) (Nityananda et al., 2015). The models were normalized such that all gave a mean response of 1 to a drifting grating at the optimal spatial and temporal frequency.

In each simulated trial, the model was presented with a 1D version of the grating used in the experiments. Energy model outputs were summed and averaged over the duration of each presentation then passed through thresh(*x*) to produce a direction judgment similar to the one made by human observers in our experiment and the psychophysics experiments of Anderson and Burr (1985). When simulating the model with noisy gratings, up to 500 presentations were repeated per noise frequency point.

In simulations of insect motion detectors, response rates were calculated as the proportion of presentations in which the direction of motion computed by the model was the same as the signal component in the stimulus. In simulations of mammalian motion detectors, we calculated detection threshold as the threshold *T* of the function thresh(*x*) that resulted in the model judging motion direction correctly in 90% of the presentations.

## Acknowledgments

• GT, VN and RR are funded by a Leverhulme Trust Research Leadership Award RL-2012-019 to JR. ISP is supported by grant PSI2014-51960-P from Ministerio de Economa y Competitividad (Spain). We are grateful to Adam Simmons for excellent insect husbandry and to Sid Henriksen for helpful comments on the manuscript.

## References

Adelson EH, Bergen JR (1985) Spatiotemporal models for the perception of motion. JOSA A 2:284–299.

Anderson SJ, Burr DC (1985) Spatial and temporal selectivity of the human motion detection system. Vision Research 25:1147–1154.

Anderson SJ, Burr DC (1989) Receptive field properties of human motion detector units inferred from spatial frequency masking. Vision Research 29:1343–1358.

Badcock DR (1984) Spatial phase or luminance profile discrimination? Vision Research 24:613–623.

Batista J, Araújo H et al. (2013) Stereoscopic depth perception using a model based on the primary visual cortex. PLOS ONE 8:e80745.

Blakemore Ct, Campbell F (1969) On the existence of neurones in the human visual system selectively sensitive to the orientation and size of retinal images. The Journal of physiology 203:237.

Borst A (2014) Neural circuits for elementary motion detection. Journal of neurogenetics 28:361–373.

Borst A, Helmstaedter M (2015) Common circuit design in fly and mammalian motion vision. Nature Neuroscience 18:1067–1076.

Brainard DH (1997) The psychophysics toolbox. Spatial Vision 10:433–436.

Buchner E (1976) Elementary movement detectors in an insect visual system. Biological Cybernetics 24:85–101.

Burge J, Geisler WS (2014) Optimal disparity estimation in natural stereo images. Journal of Vision 14:1–1.

Burr DC (1980) Sensitivity to spatial phase. Vision Research 20:391–396.

Burr DC, Ross J, Morrone MC (1986a) Smooth and sampled motion. Vision Research 26:643–652.

Burr D, Ross J, Morrone M (1986b) Seeing objects in motion. Proceedings of the Royal Society of London B: Biological Sciences 227:249–265.

Burton G (1973) Evidence for non-linear response processes in the human visual system from measurements on the thresholds of spatial beat frequencies. Vision Research 13:1211–1225.

Campbell FW, Robson J (1968) Application of Fourier analysis to the visibility of gratings. Journal of Physiology 197:551.

Carandini M (2006) What simple and complex cells compute. The Journal of Physiology 577:463–466.

Carandini M, Demb JB, Mante V, Tolhurst DJ, Dan Y, Olshausen BA, Gallant JL, Rust NC (2005) Do we know what the early visual system does? The Journal of Neuroscience 25:10577–10597.

Chen HW, Jacobson LD, Gaska JP, Pollen DA (1993) Cross-correlation analyses of nonlinear systems with spatiotemporal inputs (visual neurons). IEEE transactions on biomedical engineering 40:1102–1113.

Chichilnisky E (2001) A simple white noise analysis of neuronal light responses. Network: Computation in Neural Systems 12:199–213.

Clifford CW, Ibbotson M (2002) Fundamental mechanisms of visual motion detection: models, cells and functions. Progress in Neurobiology 68:409–437.

Daugman JG (1984) Spatial visual channels in the Fourier plane. Vision Research 24:891–910.

De Valois KK, Tootell R (1983) Spatial-frequency-specific inhibition in cat striate cortex cells. The Journal of Physiology 336:359.

Dvorak D, Srinivasan M, French A (1980) The contrast sensitivity of fly movement-detecting neurons. Vision Research 20:397407.

Emerson RC, Bergen JR, Adelson EH (1992) Directionally selective complex cells and the computation of motion energy in cat visual cortex. Vision Research 32:203–218.

Graham N, Nachmias J (1971) Detection of grating patterns containing two spatial frequencies: A comparison of single-channel and multiple-channels models. Vision Research 11:251–IN4.

Harvey LO, Gervais MJ (1978) Visual texture perception and Fourier analysis. Perception & Psychophysics 24:534–542.

Hassenstein B, Reichardt W (1956) Systemtheoretische Analyse der Zeit-, Reihenfolgen-und Vorzeichenauswertung bei der Bewegungsperzeption des Rüsselkäfers Chlorophanus. Zeitschrift für Naturforschung B 11:513–524.

Hunter I, Korenberg M (1986) The identification of nonlinear biological systems: Wiener and Hammerstein cascade models. Biological Cybernetics 55:135–144.

Kleiner M, Brainard D, Pelli D, Ingling A, Murray R, Broussard C et al. (2007) Whats new in Psychtoolbox-3. Perception 36:1.

Lawton TB (1984) The effect of phase structures on spatial phase discrimination. Vision Research 24:139–148.

Legge GE (1976) Adaptation to a spatial impulse: implications for Fourier transform models of visual processing. Vision Research 16:1407–1418.

Levinson E, Sekuler R (1975) The independence of channels in human vision selective for direction of movement. The Journal of Physiology 250:347–366.

Lu ZL, Sperling G (1995) The functional architecture of human visual motion perception. Vision Research 35:2697–2722.

Maffei L, Fiorentini A (1973) The visual cortex as a spatial frequency analyser. Vision Research 13:1255–1267.

Marr D, Hildreth E (1980) Theory of edge detection. Proceedings of the Royal Society of London B: Biological Sciences 207:187–217.

Meister M, Berry MJ (1999) The neural code of the retina. Neuron 22:435–450.

Morgan MJ (1992) Spatial filtering precedes motion detection. Nature 355:344–346.

Morrone MC, Burr D (1988) Feature detection in human vision: A phase-dependent energy model. Proceedings of the Royal Society of London B: Biological Sciences 235:221–245.

Nityananda V, Tarawneh G, Errington S, Serrano-Pedraza I, Read JC (2017) The optomotor response of the praying mantis is driven predominantly by the central visual field. Journal of Comparative Physiology A in press.

Nityananda V, Tarawneh G, Jones L, Busby N, Herbert W, Davies R, Read JC (2015) The contrast sensitivity function of the praying mantis *Sphodromantis lineola*. Journal of Comparative Physiology A 201:741–750.

OCarroll D, Laughlin S, Bidwell N, Harris R (1997) Spatiotemporal properties of motion detectors matched to low image velocities in hovering insects. Vision Research 37:34273439.

OCarroll, DC and Bidwell, NJ and Laughlin, SB and Warrant, EJ (1996) Insect motion detectors matched to visual ecology. Nature 382:6366.

Patterson RD, Nimmo-Smith I (1980) Off-frequency listening and auditory-filter asymmetry. The Journal of the Acoustical Society of America 67:229–245.

Pelli DG (1997) The VideoToolbox software for visual psychophysics: Transforming numbers into movies. Spatial Vision 10:437–442.

Pick B, Buchner E (1979) Visual movement detection under light-and dark-adaptation in the fly, *Musca domestica*. Journal of Comparative Physiology 134:45–54.

Pollen DA, Gaska JP, Jacobson LD (1988) Responses of simple and complex cells to compound sine-wave gratings. Vision Research 28:25–39.

Qian N, Andersen R, EH A (1994) Transparent motion perception as detection of unbalanced motion signals. III. Modeling. Journal of Neuroscience 14:7381–7392.

Robson J (1966) Spatial and temporal contrast-sensitivity functions of the visual system. Journal of the Optical Society of America 56:1141–1142.

Rossel S (1979) Regional differences in photoreceptor performance in the eye of the praying mantis. Journal of Comparative Physiology 131:95–112.

Sachs MB, Nachmias J, Robson JG (1971) Spatial-frequency channels in human vision. JOSA 61:1176–1186.

Snyder AW, Stavenga DG, Laughlin SB (1977) Spatial information capacity of compound eyes. Journal of Comparative Physiology 116:183–207.

Stromeyer CF, Julesz B (1972) Spatial-frequency masking in vision: Critical bands and spread of masking. Journal of the Optical Society of America 62:1221–1232.

Van Santen JP, Sperling G (1984) Temporal covariance model of human motion perception. Journal of the Optical Society of America A 1:451–473.

Van Santen JP, Sperling G (1985) Elaborated Reichardt detectors. Journal of the Optical Society of America A 2:300–321.

Zhou YX, Baker CL (1993) A processing stream in mammalian visual cortex neurons for non-Fourier responses. Science 261:98–98.

